# Multi-input drug-controlled switches of mammalian gene expression based on engineered nuclear hormone receptors

**DOI:** 10.1101/2023.02.01.526549

**Authors:** Simon Kretschmer, Nicholas Perry, Yang Zhang, Tanja Kortemme

## Abstract

Protein-based switches that respond to different inputs to regulate cellular outputs, such as gene expression, are central to synthetic biology. For increased controllability, multi-input switches that integrate several cooperating and competing signals for the regulation of a shared output are of particular interest. The nuclear hormone receptor (NHR) superfamily offers promising starting points for engineering multi-input-controlled responses to clinically approved drugs. Starting from the VgEcR/RXR pair, we demonstrate that novel (multi-)drug regulation can be achieved by exchange of the ecdysone receptor (EcR) ligand binding domain (LBD) for other human NHR-derived LBDs. For responses activated to saturation by an agonist for the first LBD, we show that outputs can be boosted by an agonist targeting the second LBD. In combination with an antagonist, output levels are tunable by up to three simultaneously present small-molecule drugs. Such high-level control validates NHRs as a versatile, engineerable platform for programming multi-drug-controlled responses.

## Introduction

Cell-based switches that regulate functional outputs in response to user-defined molecular inputs form the basis for diverse synthetic biological circuits and their biomedical applications^1, 2^. While protein-level circuits are increasingly being developed^3^, in most cases, cellular outputs are programmed through the control of target gene expression^4^. Since methods for engineering transcriptional units are well established, a diverse range of outputs can be programmed with relative ease. In contrast, there is still a limited repertoire of molecular sensors that recognize inputs, such as small molecules, and that transduce the signal to a functional response, such as gene expression. Ideally, gene outputs should be adjustable in a highly precise and bidirectional manner. Compared to single-input/single-output switches, one strategy to achieve higher-level control is through the integration of multiple molecular inputs that regulate a shared molecular output^5–13^. For example, for two cooperating signals, a first input could activate target gene expression and a second input could boost the response further, such that maximal output levels are only accessible in the simultaneous presence of both signals. Conversely, it may be desirable to tune down a previously activated response with a competing input. Ideally, to minimize the number of switch components and associated sources of noise, regulation would occur at a single promoter. This regulation would avoid the complexities of systems where important regulatory components, such as those of AND or NOR logic gates, are expressed from separate inducible promoters.

For biomedical applications of switches in engineered mammalian cells, clinically approved smallmolecule drugs represent ideal inputs because of their well-characterized pharmacodynamic, pharmacokinetic and safety profiles as well as potential synergistic effects that could be elicited with respect to disease treatment. Several drug-controlled protein switches have been developed that are based on repurposed natural proteins and/or *de novo* designed proteins^7, 8, 14–27^ However, in most of these studies, individual switches responded to a single molecular input at a time. Recently, a multi-input switch was developed based on drug-bound complexes of a ‘receiver’ protein, namely the viral protease NS3a^7^. Receiver complexes with different drugs are recognized by distinct ‘reader’ proteins, which was exploited to program graded and switchable responses^7^. While this system could enable different applications with functionalized receiver/reader combinations^7^, it was based on the competition of drug inputs for a single shared binding pocket in the receiver protein. This feature limits the type of response behaviors and does not allow for the synergistic action of two cooperating inputs that regulate a shared output. Similarly, smallmolecule inputs in this system are limited to drugs targeting a single viral protein, which restricts the input repertoire. Moreover, the viral origin of the target may present undesired immunogenic potential. To address such limitations, alternative multi-input systems that involve drug targets other than NS3a protease would be valuable. In addition, simultaneous modulation of human drug targets could be useful for synergistic therapeutic effects.

Nuclear hormone receptors (NHRs) are a protein superfamily that appears ideally suited for the design of drug-controlled protein switches. NHRs contain a ligand binding domain (LBD), a DNA binding domain (DBD) and a transcriptional activation domain (AD)^28^ (Fig. 1a). They can control target gene expression in a ligand-dependent manner as homo- or heterodimers bound to DNA-based response elements^28^. As a superfamily of 48 different LBDs in humans, they can sense a large variety of structurally and pharmacologically diverse ligands^28^. Many of the LBDs are major drug targets for different diseases, providing a large repertoire of clinically approved drugs for potential uses as switch inputs. Notably, for applications in cell therapies, repurposing a human NHR as a drug-controlled switch component could in certain cases give rise to synergies between 1) regulating engineered cells, and 2) drugging of a disease-related target. Intriguingly, multiinput-regulated responses have been observed in certain NHR heterocomplexes, in which the agonist for a first LBD activated gene expression and the agonist for a second LBD boosted expression further, while having little effect on its own^29, 30^. Similarly, both agonist and antagonist drugs are available for several NHRs, providing opportunities to both activate and tune down outputs. Lastly, it has been shown that receptor variants can be engineered that are compromised in their response to endogenous hormones but not specific drugs. A prime example for this is a tamoxifen-controlled estrogen receptor variant with decreased affinity for 17β-estradiol^31^.

**Figure 1.**
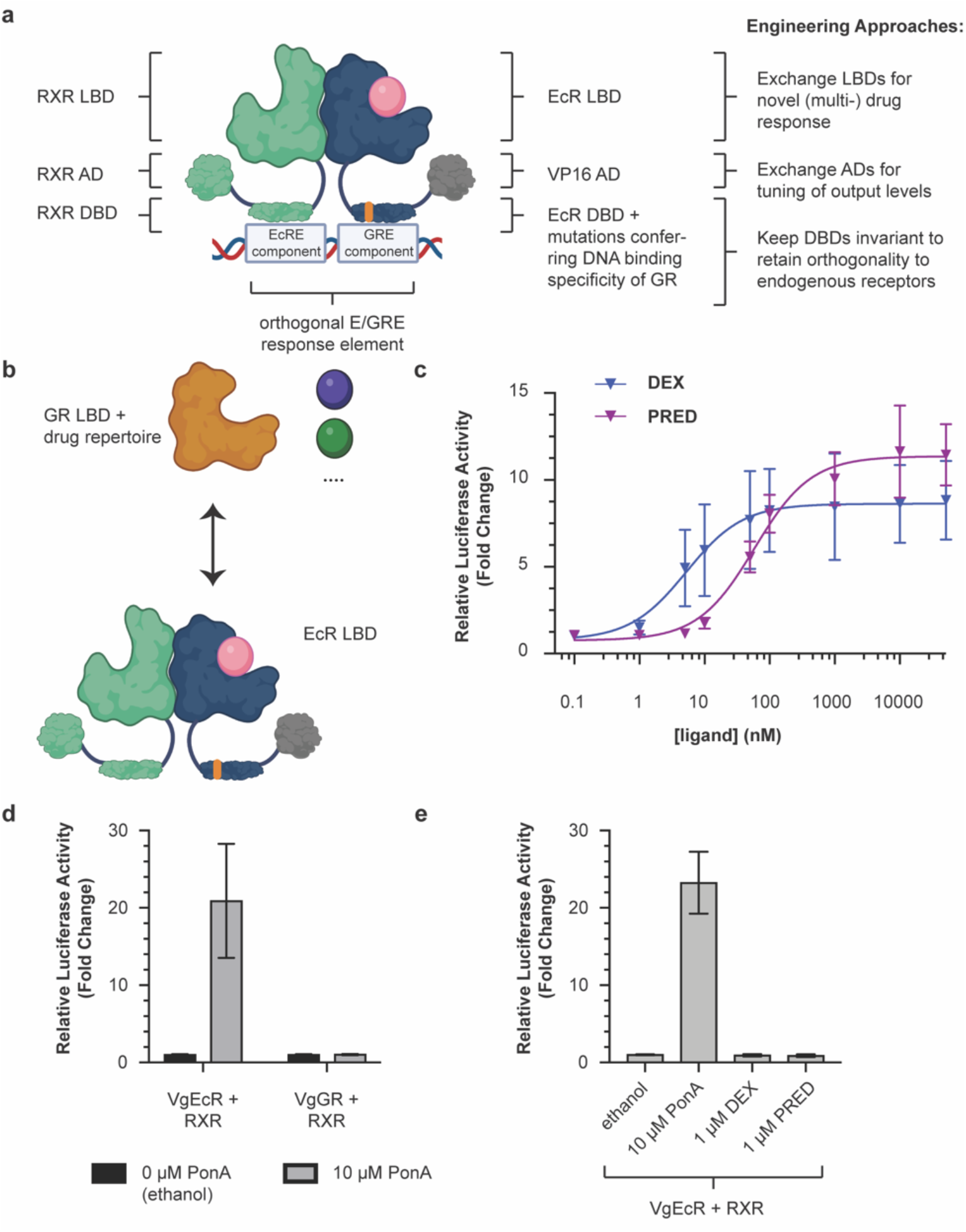
Modular exchange of the EcR LBD for the GR LBD reprograms the VgEcR scaffold to respond to GR agonists. **a)** Illustration of the original VgEcR/RXR system and engineering approaches applied to it here. **b)** Schematic of the replacement strategy for obtaining VgGR. **c)** The engineered VgGR/RXR pair activates luciferase reporter expression in response to the GR agonists dexamethasone (DEX) and prednisolone (PRED) in co-transfected HEK293 cells (n ≥ 5). Curve fits represent a three-parameter dose-response curve (Methods). **d)** The engineered VgGR/RXR system is insensitive to PonA, in contrast to the original VgEcR/RXR system (n ≥ 8). **e)** The original VgEcR/RXR pair is insensitive to DEX and PRED (n =6). Error bars represent standard deviation. Cartoons were created with BioRender.com.

Previously, NHRs or their LBDs have been used as ligand-controlled switches of gene expression in mammalian^32–35^, plant^36^ and yeast^37^ cells. A suitable starting point to engineer novel (multi-) drug-controlled responses is provided by a receptor pair^30, 34^ that was engineered for gene regulation in mammalian cells with high orthogonality, i.e. with minimal crosstalk from endogenous receptors or ligands. This pair consists of human retinoid x receptor (RXR) and a chimeric receptor based on ecdysone receptor (EcR) from *Drosophila melanogaster*, which is induced by the plant-derived ligand ponasterone A (PonA)^30, 34^ (Fig. 1a, Fig. S1). To achieve orthogonality to endogenous receptors, point mutations were introduced into the DBD of EcR to obtain the binding specificity of human glucocorticoid receptor (GR), while keeping RXR’s native DNA-binding specificity intact. This non-native combination of DBDs allows the dimer to bind to an appropriately engineered DNA sequence consisting of one half-site each of the ecdysone response element (EcRE) and the glucocorticoid response element (GRE), which is not recognized by endogenous receptors, including native GR homodimers^34^ (Fig. 1a). Besides this mutated DBD, the “VgEcR” chimera also contains the AD of VP16 from *Herpes simplex* for transcriptional activation instead of the native N-terminal domain of EcR^34^. Notably, the presence of both an EcR and an RXR agonist has been shown to produce higher output levels, when compared to an EcR agonist alone^30^, potentially enabling multi-input control for engineered derivatives of this system as well. While well-suited for orthogonal gene regulation in human cells by one or more ligands, limitations have been noted with regards to biomedical applications^38^. Of particular note, the EcR LBD responds to ligands that are not approved for clinical use and approval in the future was deemed unlikely^38^. Similarly, synergistic switch regulation and drugging of human disease targets is not possible with EcR agonists. Lastly, the LBD’s origin from insects could present undesired immunogenicity issues. To address these issues, one could envision replacing the LBD of EcR with NHR-derived LBDs from human drug targets. Yet, it is unclear how amenable the VgEcR receptor is to engineering new ligand specificities and which multi-input behaviors can arise in accordingly engineered receptors.

Here, we investigate the amenability of dimeric NHR-based systems for engineering multi-drug-controlled gene outputs. First, we reprogram the VgEcR system to respond to FDA-approved small molecules by modular exchange of the EcR LBD for human NHR-derived LBDs, while maintaining the previously engineered DBDs to retain regulation of an orthogonal target gene in human cells (Fig. 1a). Second, we demonstrate that output levels of the engineered receptor pair can be controlled by up to three simultaneously present small-molecule drugs with distinct roles in activating, boosting and counteracting target gene expression. Our results validate dimeric NHR-based systems as a versatile scaffold pair for programming multi-drug-controlled switches of gene expression in mammalian cells and motivate future use of this protein scaffold for engineering gene switches.

## Results

### Modular exchange of the EcR LBD for the GR LBD enables drug-controlled reporter expression

We first tested whether the VgEcR scaffold could be engineered to respond to agonists of human NHR ligands. We exchanged the EcR LBD with the LBD of human glucocorticoid receptor (GR), which offers a repertoire of approved drug modulators used in various disease applications (Fig. 1b). For example, dexamethasone (DEX) is used clinically to mitigate adverse effects in chimeric antigen receptor (CAR) - T cell therapies, such as “cytokine release syndrome” (CRS) or neuroinflammation^39, 40^. We did not exchange RXR’s LBD, as it can form heterodimers with many different NHRs^28^ and ligand-dependent modulation of the GR-RXR interaction has been observed^29^.

To generate a chimera between the AD-DBD portion of VgEcR and the LBD of GR, we first built a structural model of an EcR/RXR LBD dimer consisting of a homology model of VgEcR and the structure of human RXR (PDB: 3dzy). We then aligned a crystal structure of the human GR LBD (PDB: 1m2z) to the model to inform chimera design (Fig. S2a). We experimentally tested four different variants of a “VgGR” chimera (Fig. S2b) through a co-transfection assay^34, 41, 42^ in a human model cell line (Fig. S3). In particular, we transiently transfected HEK293 cells with three plasmids, namely 1) a pErv3 derivative, which constitutively expresses RXR and VgGR, 2) pEGSH-Luc, in which Firefly luciferase expression is controlled by the previously engineered VgEcR/RXR-dependent promoter, 3) pRL-CMV, which constitutively expresses Renilla luciferase as an internal control and for normalization^41^ (Fig. S3). Relative luciferase expression, as determined by the ratio of Firefly/Renilla luciferase activity and normalized to the response without ligand but appropriate solvent, was measured 20 hours after ligand addition.

In this assay, one out of the four tested VgGR chimeras, in which the EcR LBD was exchanged for the full-length GR^523-777^ LBD (VgGR-01, or simply VgGR from here on), activated reporter expression dependent on DEX in a dose-dependent manner when co-expressed with RXR (Fig. 1c, Fig. S2b,c). In addition to DEX, VgGR/RXR also responded in a dose-dependent manner to prednisolone (PRED), another drug agonist of GR (Fig. 1c). Notably, the system was more sensitive to DEX than to PRED with EC50 values of 5.2 nM for the former and 58.5 nM for the latter (Fig. 1c). As expected, the VgGR/RXR pair did not respond to PonA (Fig. 1d, Fig. S4), while the original VgEcR/RXR system did not respond to GR ligands, including DEX and PRED (Fig. 1e, S5). Taken together, drug-dependent reporter expression from the orthogonal VgEcR/RXR promoter can be reprogrammed by structure-guided modular exchange of the EcR LBD for a drug-responsive human NHR LBD.

### Cooperation of RXR and GR drug agonists can boost VgGR-induced reporter expression

Improved control over target gene expression would be achieved if maximal output levels are only accessible in the simultaneous presence of two different small molecule inputs. Therefore, we tested whether two drug inputs can cooperate in activating gene expression by the engineered VgGR/RXR system. Previously, it has been shown that RXR agonists can enhance DEX-activated reporter expression by GR from a purely GR-specific response element in COS1 cells^29^. This effect was also observed when RXR could not bind DNA by itself and was attributed to an enhanced interaction of the GR and RXR LBDs in the presence of an RXR agonist^29^. Together with the observation that RXR agonists can enhance PonA-activated expression by VgEcR^30^, this suggested that RXR agonists could also boost DEX-activated reporter expression from the orthogonal VgGR/RXR-dependent promoter.

To test this hypothesis, we performed a two-dimensional titration of DEX against the clinically approved RXR agonist drug bexarotene (BEX) (Fig. 2a,b). While DEX alone was able to activate reporter expression up to around one order of magnitude (Fig. 1c, 2b), BEX alone only activated reporter expression weakly, up to around 2-fold (Fig. 2b, Fig. S6). Strikingly, BEX was able to boost reporter expression over the entire range of tested DEX concentrations (Fig. 2b,c). In particular, BEX potentiated expression levels even at saturating concentrations of DEX (Fig. 2b,c). Thus, maximal expression levels under the tested conditions are only accessible if both DEX and BEX are present at sufficiently high concentration.

**Figure 2.**
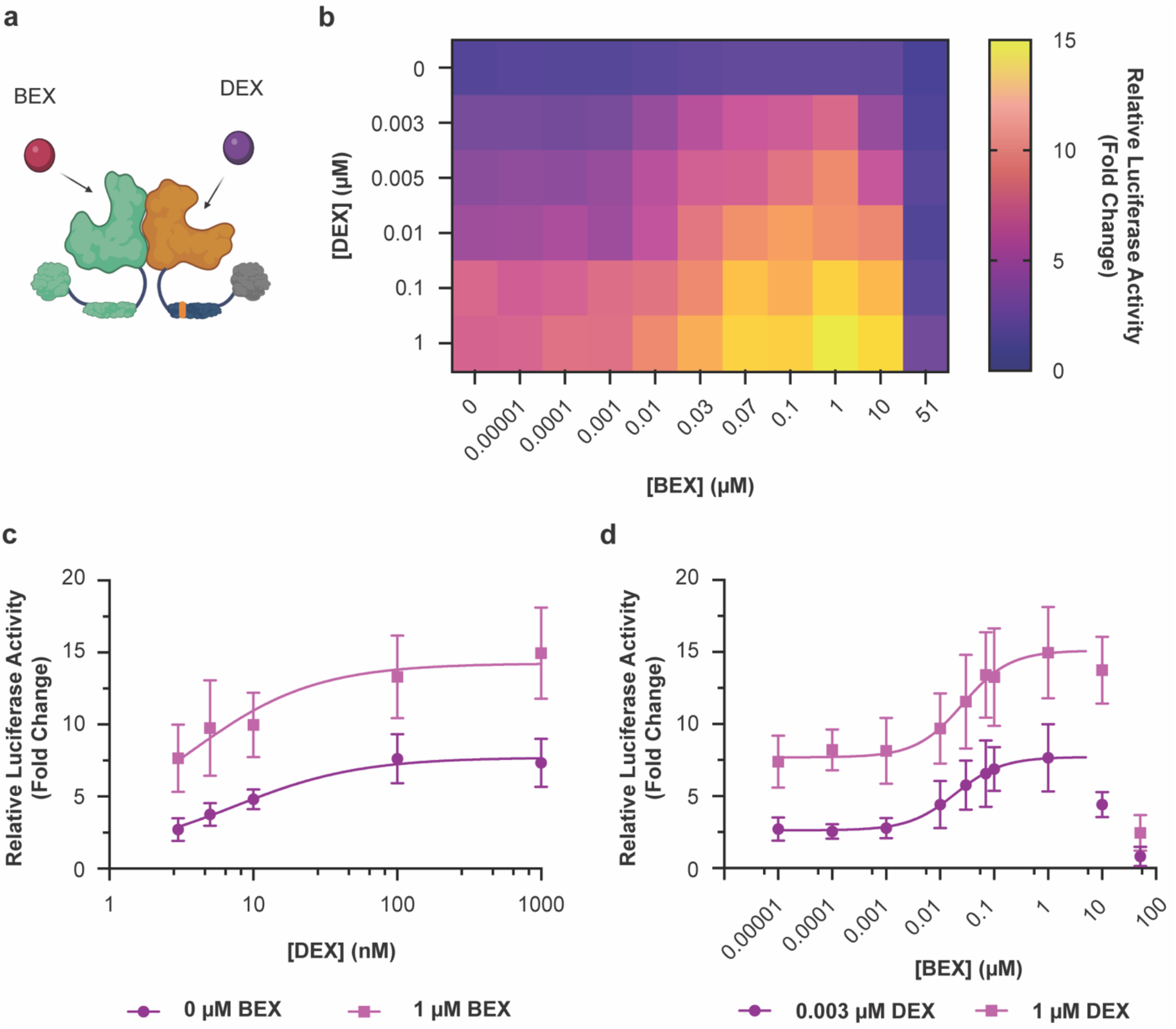
Multi-input control by cooperation of GR and RXR agonists. **a)** Illustration of the 2-input scheme for regulating the VgGR/RXR pair (created with BioRender.com). **b)** A simultaneously present RXR agonist can boost DEX-activated responses, as observed by a twodimensional titration of DEX vs. BEX (n ≥ 5). The stark drop-off in reporter expression at the highest BEX concentration is likely due to toxicity, as a similar drop-off was also observed for constitutively expressed Renilla luciferase (Fig. S7). **c)** BEX shifts DEX dose-response curves to higher output levels (n = 6). **d)** DEX shifts BEX dose-response curves to higher output levels (n = 6). Curve fits represent a three-parameter dose-response curve (Methods). The data points at 10 μM and 51 μM BEX were omitted for these curve fits due to the drop-off in reporter expression (see also Fig. S7). Error bars represent standard deviation.

When analyzing DEX and BEX dose-responses, the ligands appeared to shift each other’s outputs to higher levels rather than changing the shape of the response curves (Fig. 2c,d). Moreover, the total output in the presence of both ligands was higher than the combination of the individual ligand outputs (Fig. S6), similar to response behaviors observed in other engineered multi-input/single-output systems^10, 11, 13^. We observed a stark drop in reporter expression at the highest tested concentration of BEX (51 μM). However, as both Firefly and Renilla luciferase levels were drastically decreased under these conditions (Fig. S7), we attribute this effect to non-specific effects and/or ligand toxicity. Although this behavior defines an upper limit for BEX concentration in applications, BEX boosted reporter expression to approximate saturation over a wide window of BEX concentrations in our experiments (from 70 nM to at least 1 μM BEX) (Fig. 2b,d).

We also investigated if response parameters of the VgGR/RXR system could additionally be tuned by modular exchange of its transcriptional activation domains (Fig. S8a). In particular, we tested variants containing either the *H. simplex* VP16 domain that is also part of VgEcR^34^, the VP16-derived VP64 domain^43^ and the trimeric fusion protein VPR^44^ consisting of VP64, human p65 and Rta from *Epstein-Barr virus* (Fig. S8a). Indeed, output levels varied depending on the AD fusion (Fig. S8b). Most notably, the VPR domain substantially increased both the basal expression and output levels at saturating agonist concentrations (Fig. S8b). While the dynamic range could therefore not be increased relative to the VgGR/RXR system (Fig. S8c), this coupling of basal and maximal expression level still provides useful insights that may guide future engineering strategies, dependent on whether either tight repression without ligands or high output levels with ligand are preferred.

### Competition between GR agonist and antagonist drugs in the presence or absence of an RXR agonist drug enables 3-input control over reporter expression

In certain scenarios, it may be desirable to attenuate an activated response or tune an agonist’s functional concentration window. Toward these ends, we took advantage of the existing repertoire of clinically approved GR antagonists. In particular, we titrated DEX against RU-486 (mifepristone) to measure reporter expression at varying agonist/antagonist ratios (Fig. 3a). As anticipated, we observed maximal expression levels for high concentrations of agonist (DEX) in the absence of antagonist (RU-486) (Fig. 3b), and increasing concentrations of RU-486 resulted in a reduction of output levels (Fig. 3b,c). We note that, in the absence of DEX, RU-486 elicited an increase in reporter expression, which is consistent with this drug’s reported partial agonist effect^45^ (Fig. S9). Further, as for BEX, a non-specific decrease in luciferase expression was observed at the highest tested concentration of 51 μM RU-486 (Fig. S10).

**Figure 3.**
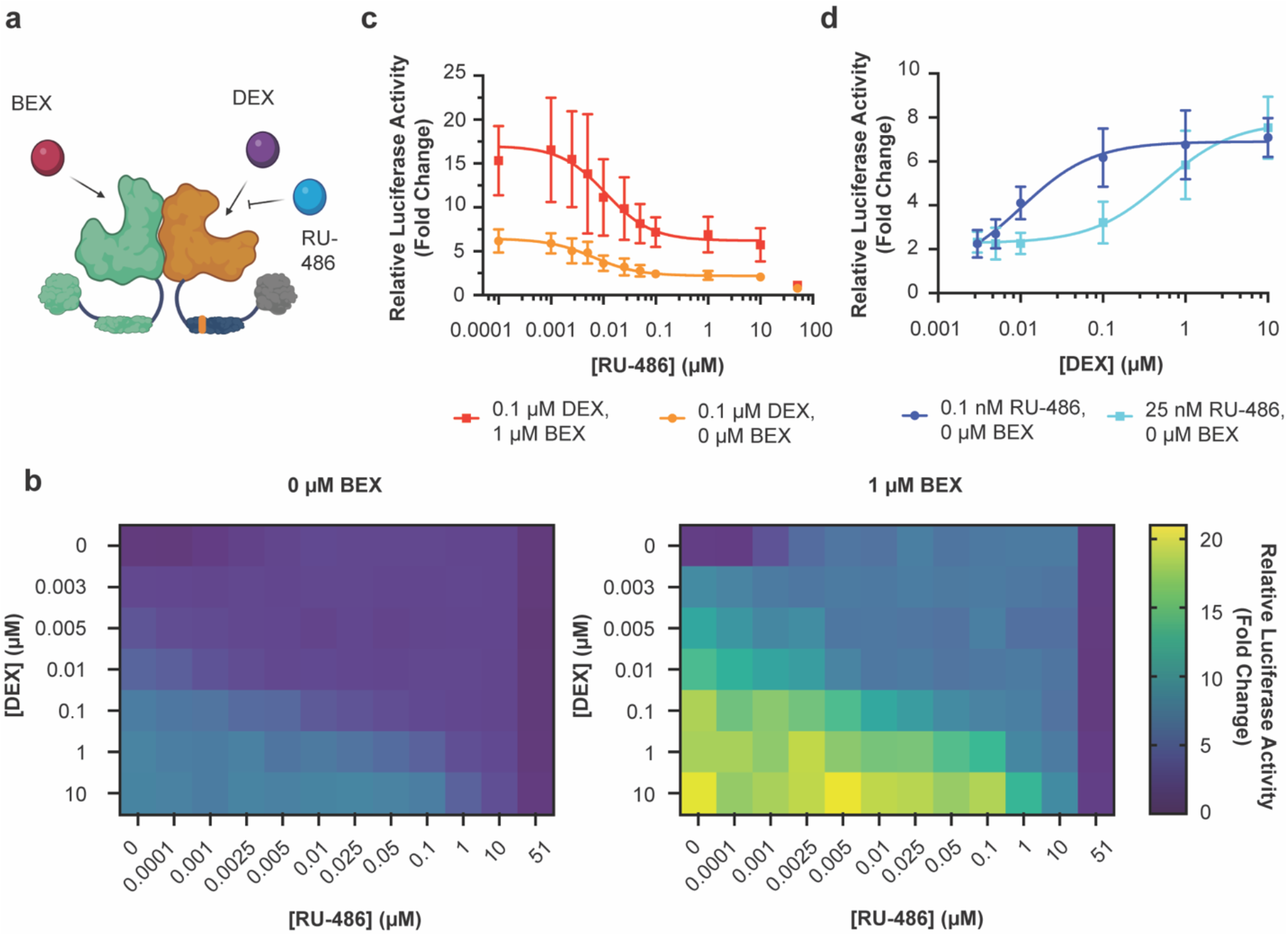
Multi-input control by simultaneous presence of GR agonists and antagonists. **a)** Illustration of the 3-input scheme for regulating the VgGR/RXR pair (created with BioRender.com). **b)** A simultaneously present GR inhibitor can attenuate DEX-activated responses, as observed by a two-dimensional titration of DEX vs. RU-486 without BEX (left) or with BEX (right) at a saturating concentration of 1 μM (Fig. 2) (n ≥ 5). **c)** BEX shifts RU-486 dose-response curves to higher output levels (n = 6). **d)** RU-486 modulates agonist sensitivity, as observed by shifting DEX dose-response to higher concentrations (n = 6). The data point at 51 μM RU-486 is omitted from these curve fits and is likely due to toxicity, as a similar drop-off was also observed for constitutively expressed Renilla luciferase (Fig. S10). Curve fits represent a three-parameter dose-response curve (Methods). Error bars represent standard deviation.

Besides enabling reduction of DEX-activated output levels, fine-tuning of both agonist and antagonist levels also modulated the agonist sensitivity, i.e. the concentration window in which a change in ligand concentration leads to a detectable change in reporter output (Fig. 3d). For example, at increasing concentrations of RU-486, higher DEX concentrations were required to both activate reporter expression and reach saturation (Fig. 3d). Lastly, we compared output reduction in the additional presence and absence of the RXR agonist BEX at a saturating concentration. While the two-dimensional response behaviors had similar properties qualitatively with or without BEX (Fig. 3b), output levels were shifted to higher levels in the presence of BEX (Fig. 3b,c). Taken together, these results demonstrate that output levels in the VgGR/RXR system can be controlled by up to three simultaneously present small-molecule ligands with distinct modulatory roles. Thereby, target gene expression can be activated by a GR agonist (DEX), boosted by an RXR agonist (BEX) and reduced by a GR antagonist (RU-486).

### Modular exchange of the EcR LBD for the estrogen receptor LBD enables reporter expression in response to additional drugs

Finally, we tested the amenability of VgEcR for modular exchange of the EcR LBD for a human NHR-derived LBD other than GR. We chose estrogen receptor (ER) due to its clinical relevance as an important drug target and correspondingly large repertoire of approved drug modulators (Fig. 4a). In addition, an ER variant (ER^T2^) has previously been engineered that displays reduced sensitivity to the endogenous ER ligand 17β-estradiol (E2), but responds to the active tamoxifen metabolite 4-hydroxytamoxifen (4OHT)^31^. Such a variant could be useful for biomedical applications due to its increased orthogonality in a physiological context and has been used in various bioengineering and synthetic biology approaches^18, 27, 31, 46^.

**Figure 4.**
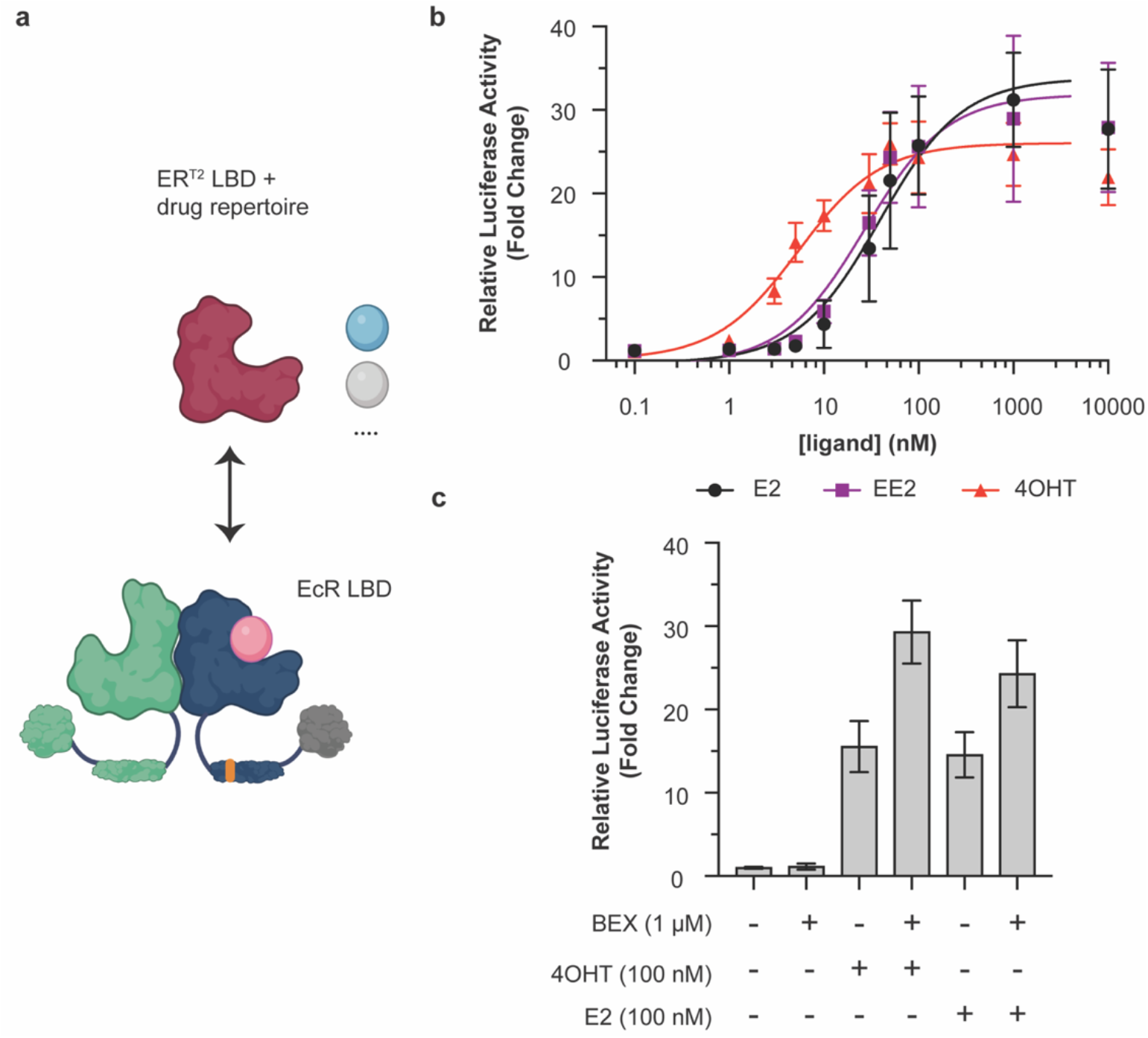
Modular exchange of the EcR LBD for the ER^T2^ LBD reprograms the VgEcR scaffold to respond to ER agonists. **a)** Schematic of the LBD replacement strategy for obtaining VgER^T2^/RXR (created with BioRender.com). **b)** The engineered VgER^T2^/RXR pair activates luciferase reporter expression in response to the ER ligands 17β-estradiol (E2), 17α-Ethynylestradiol (EE2) and 4-hydroxytamoxifen (4OHT) in co-transfected HEK293 cells (n ≥ 5). **c)** A simultaneously present RXR agonist (BEX) can boost 4OHT-activated and E2-activated responses for VgER^T2^/RXR, while having little effect on its own (n = 9). The data points at 10 μM were excluded from the curve fits due to the slight drop-off in reporter levels under these conditions. Curve fits represent a three-parameter dose-response curve (Methods). Error bars represent standard deviation.

As for GR, we generated chimeras of the AD-DBD portion of VgEcR with a small number of ER LBD variants (Fig. S11a-c). We generated two chimeras with the ERα LBD based on different fusion points in a structural alignment of the EcR and ERα LBD (PDB: 1ere). Additionally, we designed one chimera with the ER^T2^ variant, whose domain sequence was adapted from a previously reported fusion to Cre recombinase^47^. Here, we generated a fusion of the described ER^T2^ domain to VgEcR within the latter’s hinge domain between the DBD and LBD. All of the three tested chimeras were capable of activating reporter expression in response to the ER modulators E2, 17α-Ethynylestradiol (EE2) and 4OHT from the original VgEcR/RXR response element when co-expressed with RXR (Fig. 4b, S11e,f). As expected, none of the ER modulators activated reporter expression by VgEcR/RXR (Fig. S5). Interestingly, the basal and maximal output levels, dynamic range and ligand sensitivity of the VgER chimeras varied substantially between the variants (Fig. S11e,f). The ER^T2^ variant displayed the best combination of low basal activity and high fold activation (Fig. 4b, S11e,f). Additionally, in contrast to the two variants that had not been engineered for improved drug selectivity, the ER^T2^ variant was substantially more sensitive to 4OHT (EC50: 5.33 nM) than E2 (EC50: 38.76 nM) or EE2 (EC50: 26.84 nM) (Fig. 4b, Fig. S11f).

Finally, we tested if BEX could boost output levels of VgER^T2^/RXR. As for VgGR/RXR, BEX on its own had little effect on reporter expression (Fig. 4c). However, in the case of 4OHT- or E2-activated reporter expression, the simultaneous presence of BEX indeed resulted in a boosting of output levels, up to around two-fold (Fig. 4c). Curiously, we only observed BEX-mediated boosting of 4OHT- or E2-activated outputs for the engineered ER^T2^ variant, but not the ER-02 variant (Fig. S12), which lacks the mutations desensitizing the receptor against E2 and is fused to the VgEcR scaffold in a different manner (Fig. S11b,c). Nevertheless, taken together our results illustrate that two different NHR pairs (using GR or ER paired with RXR) could be engineered for (multi-)drug-controlled gene regulation from the orthogonal VgEcR/RXR promoter.

## Discussion

We have demonstrated that novel multi-drug-controlled gene regulation from an orthogonal promoter can be programmed via modular domain exchange starting from the VgEcR/RXR system. While exemplifying our strategy by replacing the EcR LBD with the GR or ER LBDs, the approach followed here should be applicable to a variety of other NHR-derived LBDs by harnessing RXR’s inherent capacity to dimerize with many other NHRs^28^. Thus, our results encourage a systematic characterization of functional dimers on the scale of the entire NHR protein superfamily, which could identify a breadth of new multi-drug-controlled switches that regulate gene outputs from the existing E/GRE response element. Conversely, the use of synthetic DBDs based on engineered zinc-finger domains^18, 48^ could allow NHR-based switches to control target expression from new DNA sites and thus enable higher-order complexity via multiplexing.

The engineered GR/RXR pair allowed for regulation of reporter expression by up to three smallmolecule drugs with distinct effects. While we characterized output levels in their simultaneous presence, the multi-input approach should be extensible to sequential drug addition schemes as well. In this way, one could envision a time-course, in which a response is first turned ON to the maximal level allowed by the activator. Subsequent addition of a booster could then effectively tune the dynamic range by increasing the achievable maximal output level in a “remote-controlled” manner. While titration of a single activating drug could, in principle, be used for step-wise increases in output levels, dual-drug-gated control of the maximal levels would be safer with regards to stochastic variations in the effective ligand concentrations within cells. Finally, if no longer required, the response could be turned OFF by an antagonist.

Notably, a simple structure-guided modular exchange strategy was sufficient to achieve responsiveness to approved drugs targeting human NHRs. This relative practical ease is an advantage over engineering workflows that involve extensive computational design and high-throughput optimization and should make the engineering of new NHR-derived gene switches more accessible in resource-limited settings. Nevertheless, we note that, while functional variants could be identified out of only a handful of tested variants for both VgGR and VgER^(T2)^, seemingly minor details in our chimera designs had notable effects. For example, truncating the GR LBD by the six most C-terminal amino acids did not result in a functional response (Fig. S2). Conversely, all three tested ER variants were functional, but with widely different sensing characteristics (Fig. S11). Such potentially unexpected sequence-function relationships indicate that the testing of multiple variants is recommended for the design of future LBD exchange variants. Moreover, AD modular exchange resulted in a simultaneous change in basal and maximal output levels (Fig. S8). To address this trade-off, LBD variants could be engineered that are conditionally destabilized in the absence of ligand, such that ligand-dependent high outputs are retained, while background levels are selectively reduced. In general, our variant-specific observations highlight certain limits to the modularity of natural protein elements and raise the question how the rationally engineered chimeras tested herein would compare to *de novo* designed protein components in terms of success rate and sensor-actuator performance.

The NHR superfamily provides a rich and diverse drug repertoire, accounting for a significant fraction (16%) of approved small-molecule drugs^49^. In addition, human NHR protein domains appear well suited for applications in cellular therapies, such as engineered immune cells^50^. In particular, certain NHRs may be synergistically targeted during switching, as they may be overexpressed in cancer. Our study represents a proof-of-principle for the protein engineering aspects and the tested compounds were largely chosen for this purpose. Nevertheless, several factors would be important to consider regarding potential biomedical applications. In principle, the 4OHT-controlled VgER^T2^/RXR pair could form the basis of an ON-switch driving the expression of a CAR within engineered T-cells applied in ER-positive breast cancer, given progress in the general application of CAR-T cells for the treatment of solid tumors^51^. On the other hand, dexamethasone’s immunosuppressive effects and its clinical use in treating CRS and neurotoxicity^39, 40^ render the VgGR/RXR pair more appropriate as an OFF-switch. In this case, one could envision controlling the expression of anti-inflammatory cytokines^52^, a pro-apoptotic caspase^22^ or a transcriptional repressor reducing CAR expression from a different promoter. While ER^T2^ was previously engineered for reduced sensitivity to endogenous hormones^31^, other receptors including GR could benefit from similar approaches regarding their ligand sensitivity. However, although tissue levels of GR ligands depend on a given physiological context, we have not observed substantial background activation for the VgGR/RXR pair compared to VgEcR/RXR (Fig. S4). As with other drug-controlled gene switches, limiting factors including the possibility of a drug’s targeting of multiple receptors^53^, potential drug-drug interactions and the therapeutic windows of drugs would have to be considered on a case-by-case basis. Moreover, for implementing gene switches in immune cells, it would be important to achieve stable integration and expression of our NHR-based components in clinically relevant cell types and to test if pharmacologically relevant absolute levels of target gene outputs can be achieved via drug induction as compared to their constitutive expression.

Taken together, we have validated the amenability of the VgEcR/RXR pair to engineer novel multi-drug-regulated gene expression, which inspires new directions from both a protein engineering and synthetic biology perspective.

## Methods

### In silico design of NHR chimeras

As no crystal structure of the VgEcR/RXR dimer is available, we built a structural model. For VgEcR, we generated a homology model using Swiss-Model^54^ with PDB structure 4nqa as template. For human RXR, we used the available PDB structure 3dzy. We then aligned the VgEcR homology model and the RXR crystal structure to an orthologous dimer structure, i.e. the PonA-bound EcR in complex with the RXR homolog Ultraspiracle (USP) from *Tribolium castaneum*, using the PyMOL Molecular Graphics System, Version 2.4.2 Schrödinger, LLC. From this model, we extracted the complex consisting of the newly oriented VgEcR/RXR LBDs as well as PonA. Lastly, we parameterized the PonA ligand and relaxed the ligand-bound dimer structure with Rosetta^55^ using coordinate constraints.

To generate VgGR, the DEX-bound human GR LBD from PDB structure 1m2z was aligned to the PonA-bound VgEcR LBD within our VgEcR/RXR LBD dimer model. Four variants were tested, in all of which the GR LBD, starting at GR residue 523, was inserted at the C-terminus of VgEcR residue 293 (Fig. S2a,b). All four variants contained the F602S mutation in GR, which was also part of the crystallized protein and is thought to increase solubility^56^. VgGR (VgGR-01) contained the full-length LBD (GR^523-777 / F602S^). VgGR-02 encompasses VgGR-01 truncated by the C-terminal loop (VgGR^523-771 / F602S^). VgGR-03 additionally contains the I628A mutation, which was described to prevent GR homodimerization (VgGR^523-771 / F602S / I628A^). VgGR-04 is a chimera of the VgGR-03 LBD with the central interface helix of VgEcR (GR^523-711 / F602S / I628A^-VgEcR^487-520^ -GR^751-771^), as observed in our alignment (Fig. S2a).

To generate VgER-01 and −02, the 17β-estradiol-bound ER LBD from PDB structure 1ere was aligned to the PonA-bound VgEcR LBD (Fig. S11a). The resulting domain orientation was reasonably well recapitulated by AlphaFold2-multimer^57^/ColabFold^58^, although a stretch at ER’s C-terminus appeared disordered (Fig. S11d). ER-01 and ER-02 differed with respect to their fusion to the VgEcR scaffold (Fig. S11c). In ER-01, the N-terminal helix of ER was retained and the N-terminus of the EcR LBD was truncated to keep as much of the ER LBD intact as possible, resulting in VgEcR^1-293^-ER^305-595^. In ER-02, the N-terminal helix of ER was truncated and the remaining LBD thus merged more seamlessly into VgEcR’s N-terminal helix, resulting in VgEcR^1-297^-ER^312-595^. For generating VgER^T2^, the final ER^T2^ sequence length was determined based on a previous successful fusion to Cre recombinase^47^. An alignment of PDB structure 3ert also appeared globally similar to the one with the unmutated ER LBD (Fig. S11a). As the functional ER^T2^ sequence within the context of the Cre fusion contained an additional N-terminal portion of ER not part of the LBD’s crystal structure, a sequence alignment was performed with Clustal Omega^59^. This alignment suggested an appropriate insertion point of the ER^T2^ sequence within the hinge domain of VgEcR connecting the DBD and LBD, resulting in VgEcR^1-268^-ER^283-594 / G400V / M543A / L544A^.

AD variants were designed through simple replacement of the VP16 sequence of VgEcR (residues 12 - 89) or the N-terminal domain of RXR (residues 2 - 99) with the tested AD sequences. Details are summarized in supplementary file “Construct_Details”.

### DNA constructs

pErv3 (GenBank: AF098284.1), pEGSH (GenBank: AF104248.3) and pEGSH-Luc were obtained as part of the “Complete Control Vector Kit” from Agilent Technologies Inc.. pRL-CMV was obtained from Promega Corporation. First, we modified pErv3, such that VgEcR and RXR (derivatives) are both tagged. While we determined an N-terminal myc tag within the VgEcR ORF, we did not identify tags in RXR. Therefore, we generated N- and C-terminal fusions of RXR with 3x-FLAG or 3-HA tags, all four of which produced a functional response to PonA when coexpressed with VgEcR (Fig. S1). Unless stated otherwise, all experiments were performed with RXR-3xFLAG (simply referred to as RXR here) or derivatives thereof. As an exception, during prototyping of GR variants, VgGR-02, VgGR-03 and VgGR-04 were co-expressed from their vectors with 3xHA-RXR. To generate the pErv3 derivatives expressing RXR-3xFLAG and RXR-3xHA, we used the Gibson Assembly^®^ master mix (NEB) with PCR products amplified from pErv3 and custom gene fragments (IDT gBlocks™) for the tags. To generate the pErv3 derivatives expressing 3xFLAG-RXR and 3xHA-RXR, restriction-based cloning using pErv3 and custom gene fragments (IDT gBlocks™) was performed. pErv3 derivatives expressing GR, ER and AD variants were obtained with PCR-amplified gene fragments (IDT gBlocks™) using restrictionbased cloning with enzymes purchased from NEB. Information on tested plasmids, including DNA coding sequences and protein sequences of VgEcR and RXR (derivatives) as well as details of their construction (enzymes, gene fragment and primer sequences) are summarized in supplementary file “Construct_Details” and tabs 1 – 3 therein.

### Small molecules

PonA was obtained from Agilent Technologies Inc./Invitrogen. Dexamethasone, prednisolone, E2, EE2 and 4OHT were obtained from MilliporeSigma. Bexarotene was obtained from Thermo Scientific. RU-486 was obtained from Torcis Bioscience. Stock solutions in ethanol were prepared for all compounds at 100x of their final assay concentrations.

### Transient transfection of HEK293 cells and luciferase reporter assays

The luciferase reporter assay was adapted from previous protocols^41, 42^. First, HEK293 cells (obtained from the UCSF Cell Culture Core Facility) were seeded in a sterile, tissue culture treated, white, flat-bottom 96-well microplate (Corning^®^, product number 3917) at a density of 3 * 10^4^ cells per well in 100 μL DMEM medium supplemented with 10 % FBS (Thermo Fisher Scientific). After incubation at 37 °C and 5% CO2, cells were transiently transfected on the next day using lipofectamine™ 2000 transfection reagent (Invitrogen, Thermo Fisher Scientific), according to the manufacturer’s instructions. Transfection mixes were prepared, such that addition of 10 μL of DNA:lipid complexes in DMEM resulted in 24 ng pErv3 derivative, 95 ng pEGSH-Luc, 2 ng pRL-CMV and 0.4 μL lipofectamine™ 2000 reagent per well. Cells in background wells were transfected with 95 ng empty pEGSH vector. Upon incubation for approximately 24 h, the media was exchanged for 75 μL DMEM medium supplemented with 10 % FBS per well. On the following day, small molecules from 100x stock solutions, or ethanol for measuring basal expression, were added to the cells. Finally, after incubation for 20 h, Firefly and Renilla luciferase activity were measured in a Synergy H1 microplate reader (BioTek) using the Dual-Glo^®^ Luciferase Assay System (Promega) according to the manufacturer’s instructions.

### Data Analysis

For both the Firefly and Renilla luciferase activities in a given sample well, an average of the reads from the background wells was subtracted. The Firefly/Renilla Luc Ratio was then calculated from these background-corrected levels. The relative luciferase activity (fold change) represents the Firefly/Renilla Luc Ratio in the presence of a small molecule, normalized by the Firefly/Renilla Luc ratio in the absence of small molecules but with ethanol. To calculate this property, we divided the Firefly/Renilla Luc Ratio of every drug-containing sample well by an average of the Firefly/Renilla Luc Ratio for same-day replicate wells without drug. The plotted values were then obtained by averaging relative luciferase activities from same/multi-day replicates.

Dose-response curves, heat maps and bar plots were generated with Prism 9 (GraphPad). Nonlinear curve fits were obtained with the built-in “Agonist vs. response (three parameters)” [Y=Bottom + X*(Top-Bottom)/(EC50 + X)] and “Inhibitor vs. response (three parameters)” [Y=Bottom + (Top-Bottom)/(1+(X/IC50))] equations. EC50 estimates were obtained from such curve fits to relative luciferase activity data.

## Supporting information

Supplementary file "Construct_Details"

## Supporting Information

Figures S1 – S12

Supplementary file “Construct_Details”

## Abbreviations

EcR: ecdysone receptor;
RXR: retinoid x receptor;
GR: glucocorticoid receptor;
LBD: ligand binding domain;
DBD: DNA binding domain;
AD: activation domain;
NHR: nuclear hormone receptor;
FDA: U.S. Food and Drug Administration;
PonA: ponasterone A;
DEX: dexamethasone;
PRED: prednisolone;
BEX: bexarotene;
E2: 17β-estradiol;
EE2: 17α-Ethynylestradiol;
40HT: 4-hydroxytamoxifen;
E/GRE: Ecdysone/Glucocorticoid response element;
USP: Ultraspiracle

## Author Information

**Corresponding author** * Simon Kretschmer, E-Mail

## Author Contributions

S.K. and T.K. conceived the project. S.K. performed *in silico* chimera design, cloning, mammalian cell reporter experiments and data analysis. S.K. and N.P. built the structural model of the VgEcR/RXR dimer. N.P. performed cloning and preliminary functional characterization of tagged RXR variants. Y.Z. provided technical help with initial cell culture experiments. S.K. wrote the manuscript draft with contributions from T.K. and N.P..

## Acknowledgment

We thank members of the Kortemme lab as well as Prof. Kole Roybal (UCSF) for helpful suggestions and discussions. This work was supported by the U.S. National Institutes of Health (NIH) under Grants R01GM110089 and R35GM145236 to T.K.. S.K. was supported by a Postdoctoral Independent Research Grant of the UCSF “Program for Breakthrough Biomedical Research” (PBBR), which is partially funded by the Sandler foundation, and a Li foundation endowed fellowship. Tanja Kortemme is a Chan Zuckerberg Biohub Senior Investigator.

## Notes

### Competing Interest Statement

The authors have declared no competing interest.

