## Supporting Information for "Multi-input drug-controlled switches of mammalian gene expression based on engineered nuclear hormone receptors"

**This PDF file includes:**

**Figs S1 to S12**

### Supplementary Figures

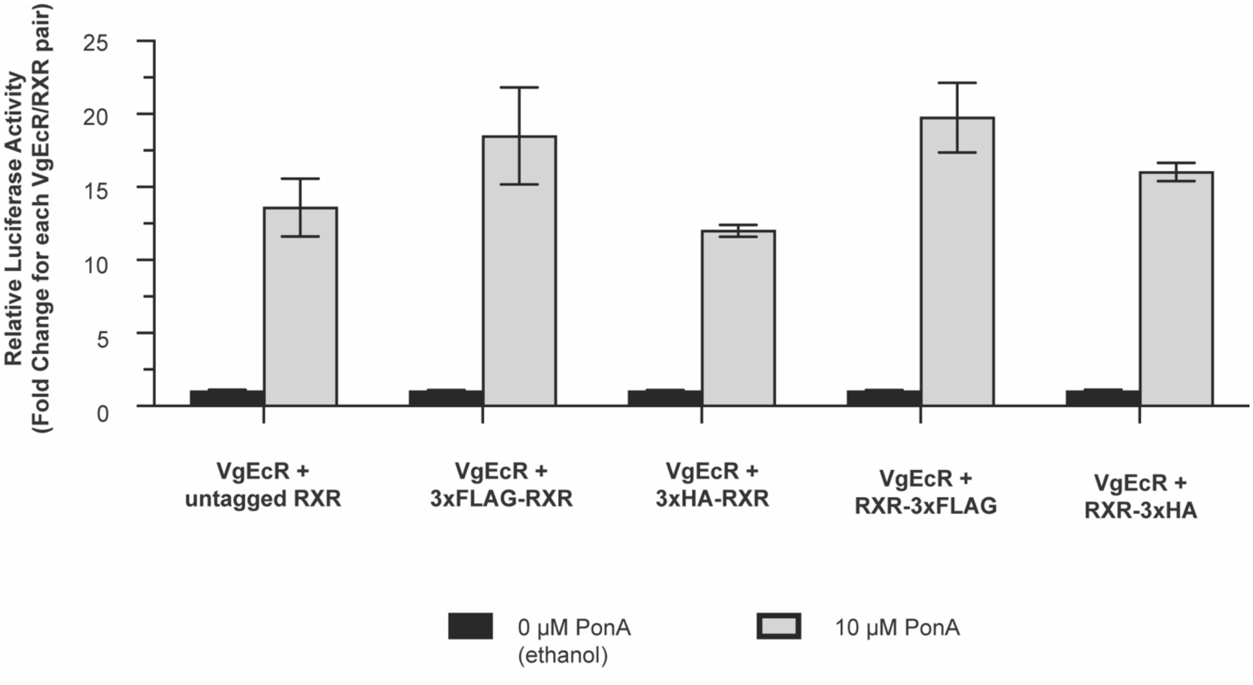


**Figure S1.** Functional characterization of N- and C-terminally tagged RXR variants. 3x repeats of FLAG (DYKDDDDK) or HA (YPYDVPDYA) tags were fused via $G_4S$ linkers to the N- or C-terminus of RXR. All RXR variants induce reporter expression in response to PonA, when co-expressed with VgEcR from a pErv3 derivative in HEK293 cells (n = 5). Error bars represent standard deviation.

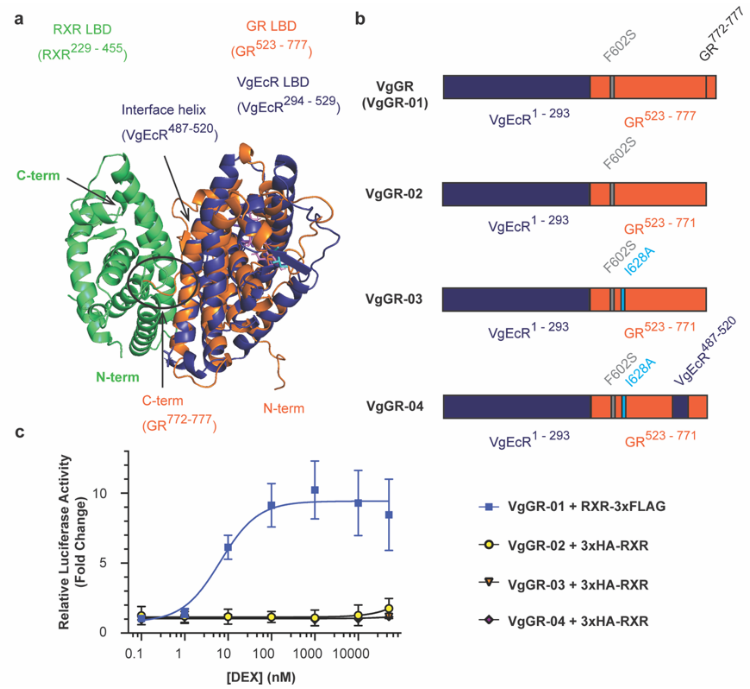


**Figure S2.** Initial prototyping of VgGR chimeras identified one functional variant out of four tested. **a)** Manual alignment of the DEX-bound GR LBD (GR$^{523\text{-}777\,/\,F602S}$) to the EcR portion of our custom-built model (see Methods) of the VgEcR/RXR LBDs. Colors: RXR, limegreen; VgEcR, deepblue; GR, orange. **b)** Construct schematics; All variants contain the solubility-enhancing F602S mutation that was also present in the crystal structure[56]. As the alignment indicated that the 6 most C-terminal amino acids (circled in **a**) would clash with RXR, these residues were removed by truncation in constructs VgGR-02, -03 and -04. The VgGR-03 and VgGR-04 constructs additionally contained a mutation of I628 (cyan sticks) to alanine, which had been described to decrease GR homodimerization[56]. VgGR-04 is a chimera based on VgGR-03, in which GR's central interface helix in the alignment is exchanged with the corresponding residues of EcR. **c)** While VgGR-01 activates reporter expression in response to DEX when co-expressed with RXR-3xFLAG, VgGR-02, -03 and -04 were insensitive to DEX when co-expressed with 3xHA-RXR (n = 3). This experiment hence identified one functional variant, while suggesting that the alignment in Fig. 1a is inaccurate. One explanation could be that the C-terminal loop in GR adopts a different conformation that does not clash with RXR. Functionality of pErv3 derivatives expressing tagged RXR variants is shown in Fig. S1. The data for VgGR-01 were collected in a separate set of experiments as the data in Fig. 1c within the main text. Curve fits represent a three-parameter dose-response curve (Methods). Error bars represent standard deviation.

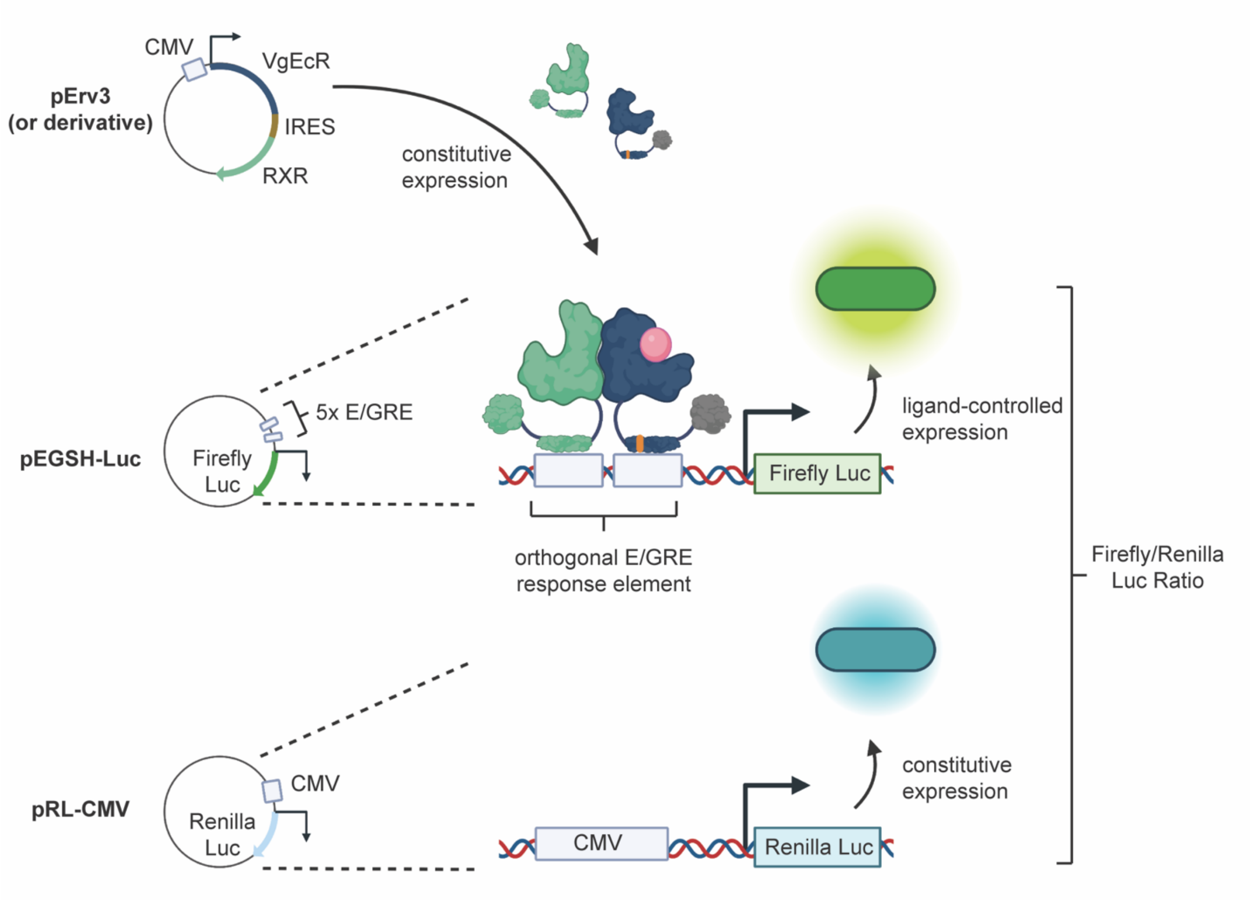


**Figure S3.** Schematic of the co-transfection assay performed to assess chimera functionality in HEK293 cells. In pErv3, a CMV promoter drives the expression of VgEcR and RXR, whose coding sequences are separated by an internal ribosome entry site (IRES) sequence. In pEGSH-Luc, the expression of firefly luciferase is controlled by ligand binding to VgEcR/RXR. Five copies of the orthogonal E/GRE response element upstream of three SP1 binding sites and a minimal heat shock promoter each bind one VgEcR/RXR pair. To account for variability due to factors like different cell numbers or transfection efficiencies between samples, Renilla luciferase is constitutively co-expressed from a CMV promoter on pRL-CMV. Gene switch induction is assessed by calculating the ratio of Firefly/Renilla luciferase activity for each condition. Created with BioRender.com.

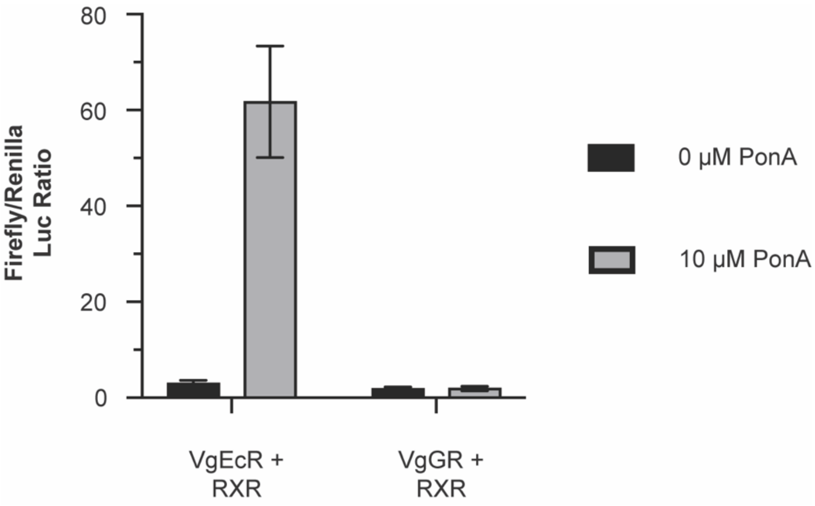


**Figure S4.** In the absence of any added ligands, the VgGR/RXR pair has comparable background activity to the original VgEcR/RXR pair, as evident when comparing the Firefly/Renilla luciferase activity ratios without normalization to the level at 0 µM PonA ($n \geq 8$). Data shown are the same as in Fig. 1d, which shows normalized relative luciferase activity . Error bars represent standard deviation.

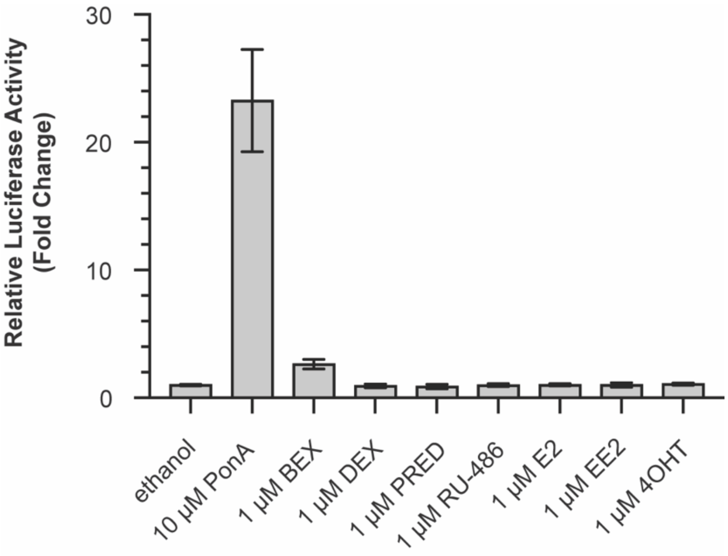


**Figure S5.** The original VgEcR/RXR system activates luciferase reporter expression in co-transfected HEK293 cells in response to the EcR agonist PonA, but not ligands of ER or GR. The RXR agonist BEX exhibits a weak stimulatory effect, as also observed for the engineered VgGR/RXR system in the absence of GR agonist (Fig. 2, Fig. S6). All ligands were added at concentrations that produced saturating effects for their respective target receptors (1µM or 10µM as indicated) (n = 6); For ethanol, PonA, DEX and PRED, the same data as in Fig. 1e are shown, which were collected in the same set of experiments with the other ligands. Error bars represent standard deviation. PonA: ponasterone A, BEX: bexarotene, DEX: dexamethasone, PRED: prednisolone, RU-486: mifepristone, E2: 17β-estradiol; EE2: 17α-Ethynylestradiol; 4OHT: 4-hydroxytamoxifen.

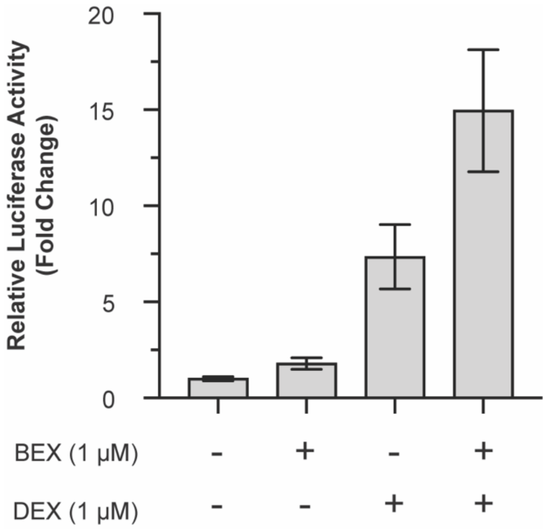


**Figure S6.** BEX boosts DEX-activated reporter expression and the total output is higher than the combined individual outputs. In particular, the difference in DEX-activated reporter expression in the presence and absence of BEX is substantially higher than output levels achieved with BEX alone (n = 6; same data as in Fig. 2). Error bars represent standard deviation.

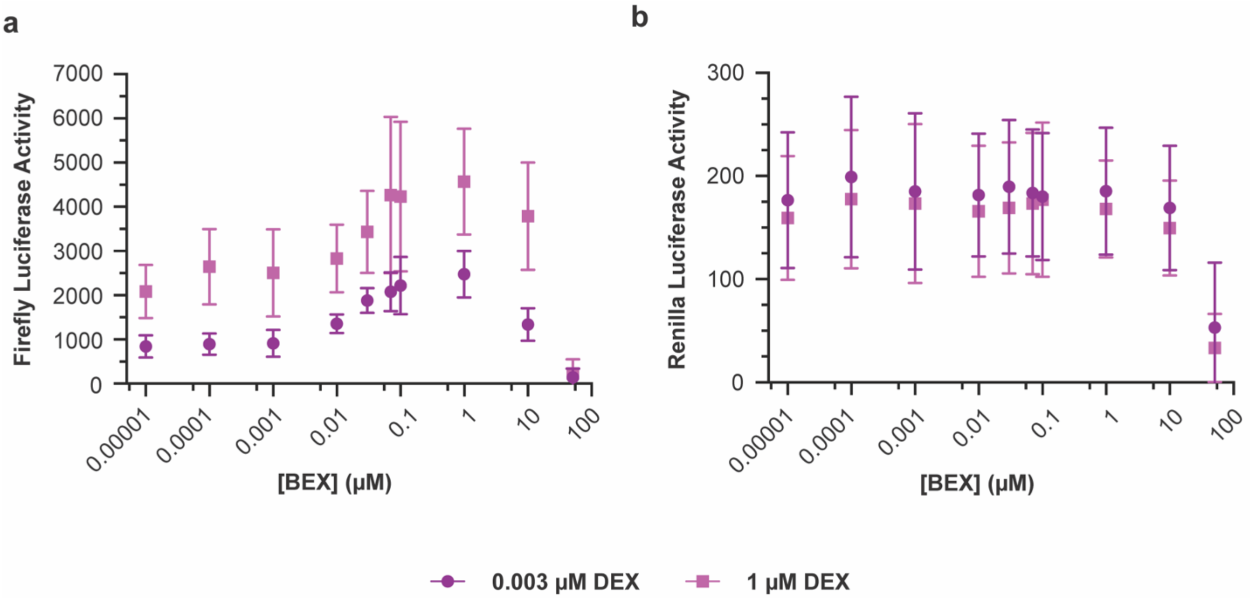


**Figure S7.** At the tested maximal concentration of 51 µM, BEX resulted in a stark drop-off in luciferase activity for both Firefly (**a**) and Renilla (**b**) luciferases, indicating non-specific effects and/or toxicity under these conditions. Data shown are the same as in Fig. 2d, which shows normalized relative luciferase activity. Error bars represent standard deviation (n = 6).

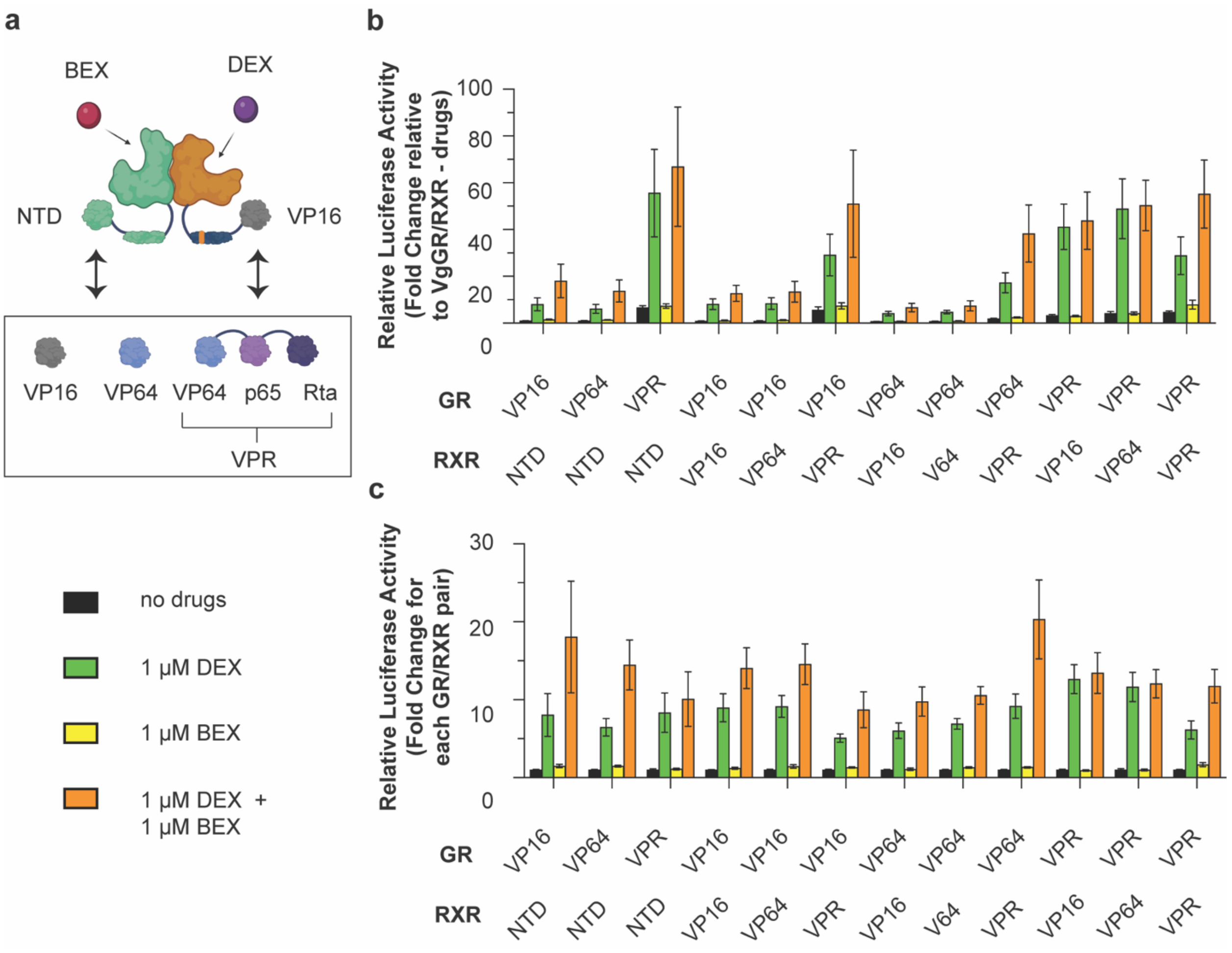


**Figure S8.** Modular exchange of transcriptional activation domains in VgGR/RXR tunes output levels. **a)** Schematic of the AD exchange strategy that was applied to the N-terminal domains (NTDs) of both RXR (RXR NTD) and VgGR (VP16) (created with BioRender.com). **b)** Normalization of reporter levels to the “no drugs” condition of the VgGR/RXR starting pair shows that AD exchange tunes the basal and maximal output levels in a coupled manner (n ≥ 5). **c)** Normalization of reporter levels to the “no drugs” condition of each individual GR/RXR pair highlights that the dynamic range was not increased relative to the VgGR/RXR starting pair (n ≥ 5). Error bars represent standard deviation.

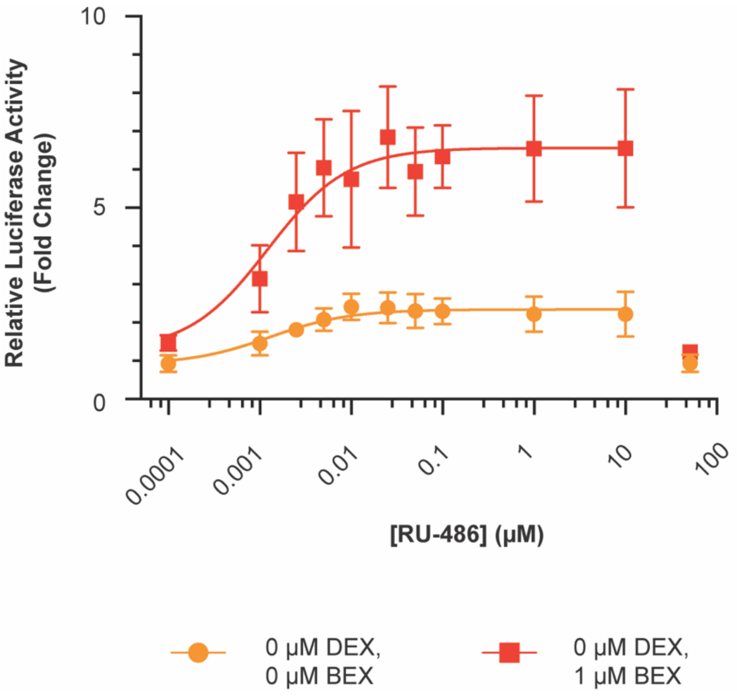


**Figure S9.** In the absence of DEX, RU-486 displays a partial agonist effect, in contrast to its antagonist activity in the presence of DEX (Fig. 3) (n = 6; same data as in Fig. 3). The data points at 51 µM RU-486 were omitted for these curve fits and are likely due to toxicity, as a similar drop-off was also observed for constitutively expressed Renilla luciferase (Fig. S10). Curve fits represent a three-parameter dose-response curve (Methods). Error bars represent standard deviation.

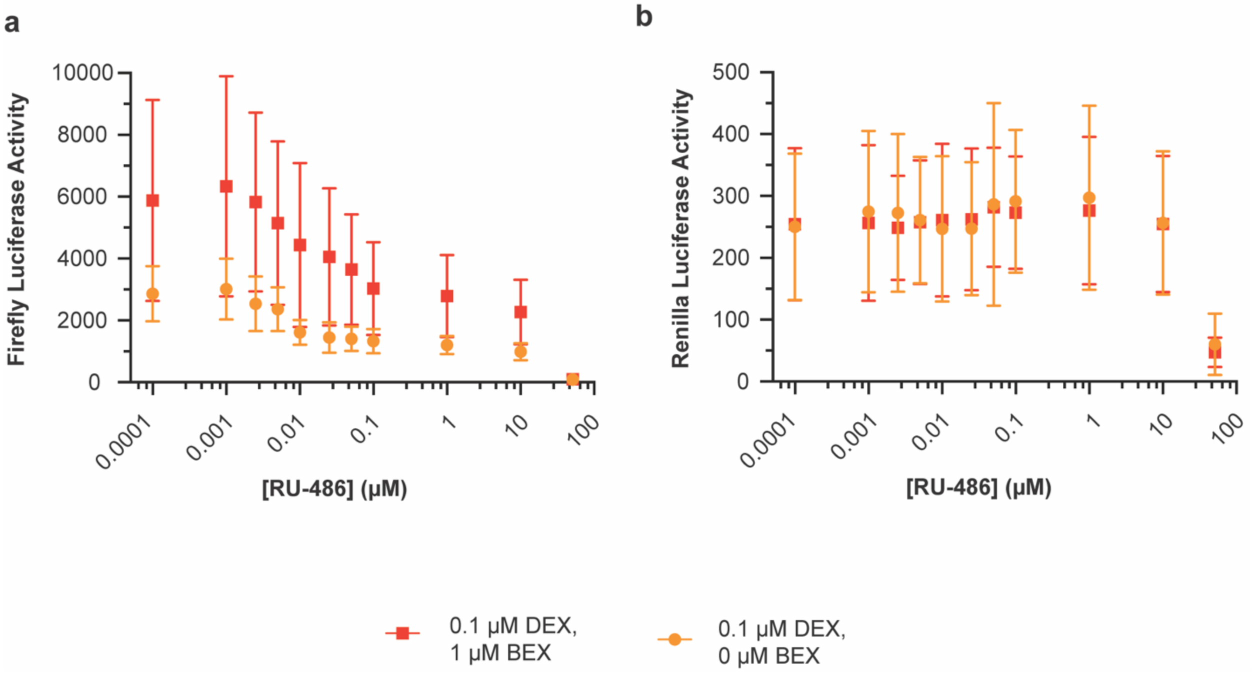


**Figure S10.** At the tested maximal concentration of 51 µM, RU-486 resulted in a stark drop-off in luciferase activity for both Firefly (**a**) and Renilla (**b**) luciferases, indicating non-specific effects and/or toxicity under these conditions. The plots show the same data as in Fig. 3d, which shows normalized relative luciferase activity (n = 6). Error bars represent standard deviation.

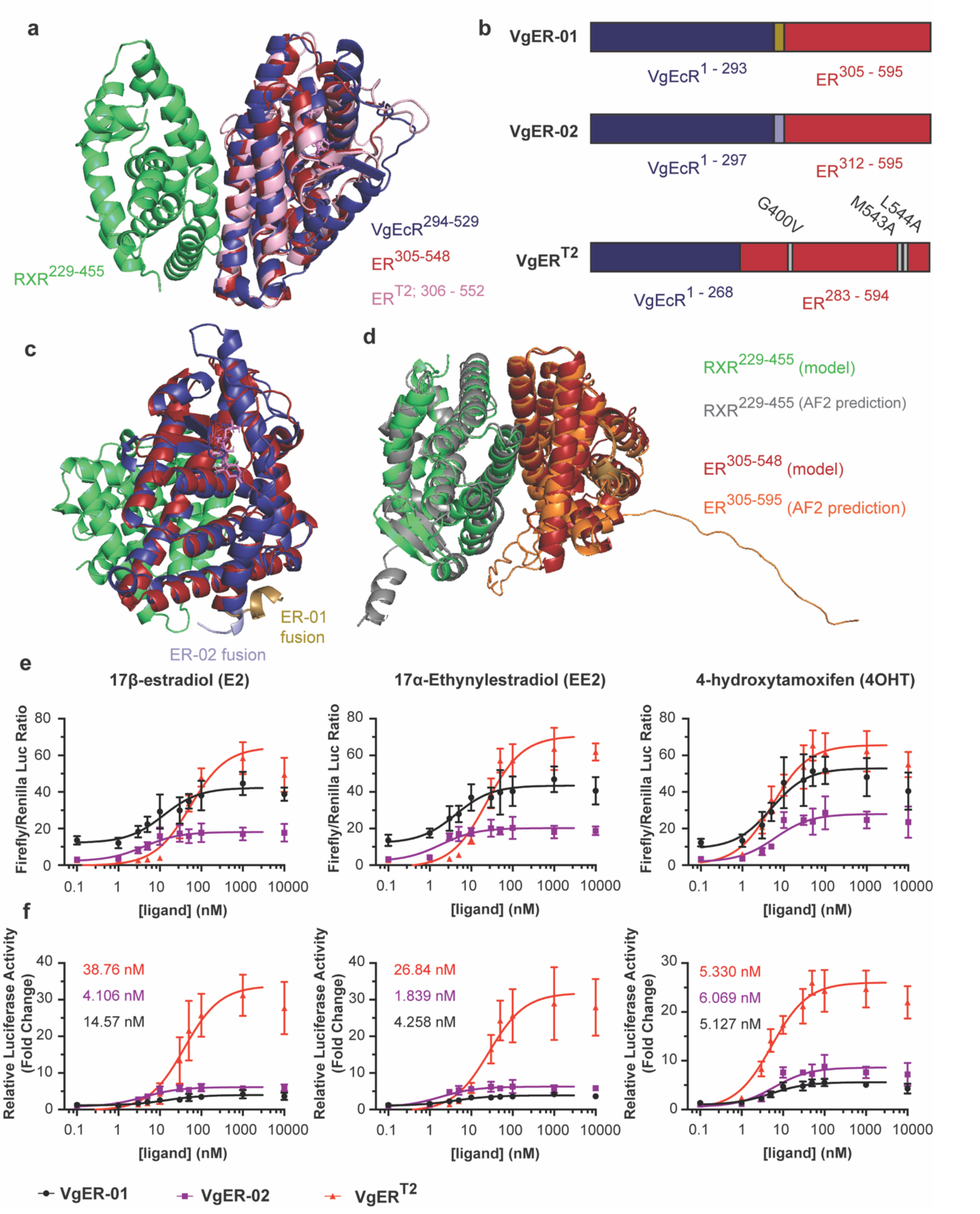


**Figure S11.** Molecular details of VgER chimeras affect sense-response parameters for different ligands. **a)** Alignment of the ER (residues 305 – 548; PDB: 1ere; firebrick) and $ER^{T2}$ (residues 306 – 552; PDB: 3ert; lightpink) LBD crystal structures to the EcR LBD portion of our custom-built model of the VgEcR (residues 294 – 529; deepblue)/RXR (residues 229 – 455; limegreen) LBDs. **b)** Schematics of the tested VgER constructs. **c)** For the VgER-01 chimera, VgEcR was truncated post-residue 293 (deleting the lightblue-colored fragment) and ER inserted starting from ER residue 305 (keeping the sand-colored fragment). In VgER-02, VgEcR was truncated post-residue 297 (keeping the lightblue-colored fragment) and ER inserted starting from ER residue 312 (deleting the sand-colored fragment). The remaining colors are identical to panel a. **d)** At the LBD interface, an AlphaFold Multimer V2 prediction of the RXR (residues 229 – 455; grey)/ER (residues 305 – 595; tvorange) complex showed reasonably good agreement with our alignment-based model (RMSD = 1.346 Å as determined with PyMOL; colors as above). **e)** Firefly/Renilla luciferase activity ratios for VgER/RXR pairs highlight differences in the basal and maximal signal for the three VgER chimeras. **f)** Relative Luciferase activities, normalized to 0 μM ligand, highlight differences in the fold change for the three VgER chimeras ($n \geq 5$). Numbers in the graphs correspond to EC50 values. The data points at 10 μM ligand were excluded from the curve fits. The data in panels e and f for $VgER^{T2}$ are the same as that plotted in Fig. 4b. Curve fits represent a three-parameter dose-response curve (Methods). Error bars represent standard deviation.

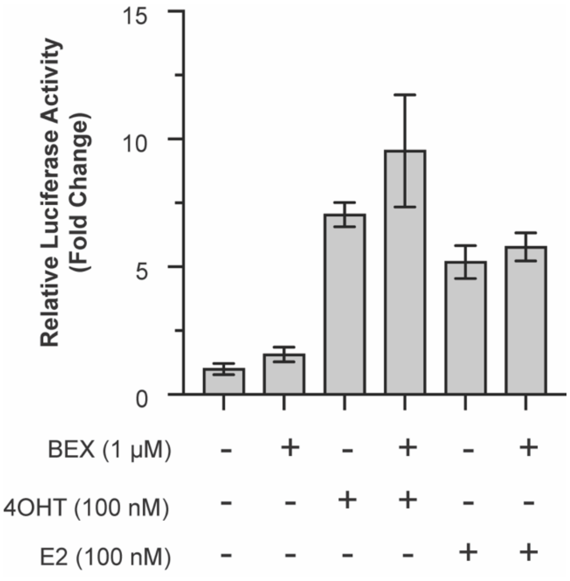


**Figure S12.** For VgER-02/RXR, there was no notable boosting of 4OHT-activated and E2-activated responses by a simultaneously present RXR agonist (BEX) (n = 6).
